# Modeling the carbon-dioxide response function in fMRI under task and resting-state conditions

**DOI:** 10.1101/2022.07.04.498727

**Authors:** Seyedmohammad Shams, Prokopis Prokopiou, Azin Esmaelbeigi, Georgios D. Mitsis, J. Jean Chen

**Author notes:** Corresponding author: Jean Chen.

## Abstract

Conventionally, cerebrovascular reactivity (CVR) is estimated as the amplitude of the hemodynamic response to vascular stimuli. While the CVR amplitude has established clinical utility, the temporal characteristics of CVR have been increasingly explored and may yield even more pathology-sensitive parameters. This work is motivated by the current need to evaluate the feasibility of dCVR modeling in various noise conditions. In this work, we present a comparison of several recently published model-based deconvolution approaches for estimating *h*(*t*), including maximum a posterior likelihood (MAP), inverse logit (IL), canonical correlation analysis (CCA), and basis expansion (using Gamma and Laguerre basis sets). To aid the comparison, we devised a novel simulation framework that allowed us to target a wide range of SNRs, ranging from 10 to −7 dB, representative of both task and resting-state CO_2_ changes. In addition, we built ground-truth *h*(*t*) into our simulation framework, overcoming the practical limitation that the true *h*(*t*) is unknown in methodological evaluations. Moreover, to best represent realistic noise found in fMRI scans, we extracted it from in-vivo resting-state scans. Furthermore, we introduce a simple optimization of the CCA method (CCA_opt_) and compare its performance to these existing methods. Our findings suggest that model-based methods can reasonably estimate dCVR even amidst high noise, and in a manner that is largely independent of the underlying model assumptions for each method. We also provide a quantitative basis for making methodological choices, based on the desired dCVR parameters, the estimation accuracy and computation time. The BEL method provided the highest accuracy and robustness, followed by the CCA_opt_ and IL methods. Of the three, the CCA_opt_ method required the lowest computational time. These findings lay the foundation for wider adoption of dCVR estimation in CVR mapping.

## Introduction

Cerebrovascular reactivity (CVR) refers to a vasodilatory or constrictive reaction of a blood vessel to a vasoactive stimulus. CVR has well-established prognostic value for cerebrovascular diseases (Glodzik et al., 2013; Pillai and Mikulis, 2015; Zhao et al., 2021), is commonly measured with the help of carbon dioxide (CO_2_) variations. CO_2_ is a potent vasodilator, and its action on cerebral blood flow (CBF) has been well documented (Battisti-Charbonney et al., 2011; Nowak-Flück et al., 2018). CO_2_-induced CBF changes in the middle cerebral artery have been measured using transcranial Doppler ultrasound (TCD) (Pinto et al., 2020). However, to achieve whole brain coverage while maintaining the monitoring of dynamic CBF changes, functional MRI (fMRI) methods are now increasingly relied upon. The fMRI method of choice in this context is the blood-oxygenation level dependent (BOLD) signal, which reflects the vascular response to CO_2_ through a non-linear relationship with CBF (Davis et al., 1998; Halani et al., 2015; Hoge et al., 1999). The measurement of the BOLD fMRI signal in response to external CO_2_ stimuli resulted in the earliest clinical application of fMRI-based CVR mapping (Blockley et al., 2017; Chen, 2018; Fierstra et al., 2013), where CVR is measured simply as the ratio of the BOLD percent response to the CO_2_ change in mmHg, which we shall refer to as “static CVR”. Quantitative static CVR, which has largely been estimated as the gradient of changes in the BOLD signal with changing end-tidal CO_2_ (PETCO_2_) values, has found promising applications, as was summarized in numerous reviews (Glodzik et al., 2013; Peng et al., 2018; Pillai and Mikulis, 2015; Pinto et al., 2020).

Recently, the temporal characteristics of CVR have been increasingly explored (Duffin et al., 2015), yielding even more promising results, leading to the term dynamic CVR (dCVR) (Prokopiou et al., 2019). dCVR provides not only CVR amplitude but also its shape, the latter being found to differ between young and older controls (West et al., 2019). Moreover, recent work showed that the BOLD timing parameter was found to better delineate between those with mild-cognitive deficit (MCI) and Alzheimer’s disease (AD) (Holmes et al., 2020). Static CVR can be readily obtained as the area under the dCVR function, representing the “steady-state” response to a step PETCO_2_ change. Additionally, specific timing parameters can also be determined. Examples include the time to peak and time to recover, which can reflect the vasodilatory and vasoconstrictive ability as well as the associated transit delays, and vary with vascular tone (Halani et al., 2015). Moreover, the width of the response function at the half-maximum point is a common shape parameter that reflects a combination of the onset and recovery responsiveness of the vasculature (Leoni et al., 2008).

The idea of modeling the BOLD response to PETCO_2_ coincided with the goal to map CO_2_-related physiological contributions in resting-state fMRI (Golestani et al., 2016, 2015). Modeling of dCVR can play a key role for physiological correction in resting-state functional-connectivity mapping (Chang and Glover, 2009; Golestani et al., 2015). The resting-state CO_2_ influence also presents a valuable opportunity to CVR (J. J. Chen et al., 2021; Jahanian et al., 2017; Liu et al., 2017). Using deconvolution based on division in the Fourier domain, Duffin et al. found the transfer function relating a block CO_2_ stimulus with the BOLD signal, whereby the phase of the transfer function could be converted to a time delay parameter (Duffin et al., 2015). An extension of this approach is the use of the Wiener filter, which was recently demonstrated for estimating the hemodynamic response to changes in neuronal local-field potential (Wu et al., 2021). However, due to the low signal-to-noise (SNR) conditions in rs-fMRI, deconvolution of resting CO_2_ fluctuations from the fMRI signal is non-trivial, and the methods that have been found adequate for estimating the neuronal or CO_2_-stimulus response may not be appropriate. Atwi et al. used a non-parametric singular-value decomposition (SVD) approach to model dCVR in both task and resting-state fMRI (Atwi et al., 2019). In the SVD method, which is well established for response-function modeling in dynamic susceptibility contrast MRI (Chen et al., 2005; Ostergaard et al., 1996; Wu et al., 2003), noise contributions are controlled by thresholding the eigenvalues of the diagonal matrix. This effectively reduces the rank of the decomposition, which is by definition analogous to truncating the frequency spectrum of the signal based on spectral power, and leads to oscillations in the resulting dCVR estimate. These oscillations can make it more challenging to obtain an accurate determination of response onset and offset times.

To overcome the uncertainties associated with the low SNR and the fact that the CO2 response can overlap in frequency with the neuronal hemodynamic response, model-based methods are generally more robust (Chen et al., 2005), but with the caveat that their performance may depend on the underlying model structure. To this point, there is currently no systematic demonstration and evaluation of deconvolution methods suited to estimating the dCVR in both high- and low-SNR conditions for various assumed dCVR models.

In this context, the aim of this work is to evaluate various dCVR modeling approaches, and in doing so, propose a unified approach to quantify the dCVR under various SNR conditions with the least bias by underlying model assumptions. We took a simulation-based approach in which we know the true SNR and the true dCVR. We implemented a total of five current modelbased deconvolution techniques, including: (1) the maximum a-posteriori (MAP) method using a Gaussian basis (Chang et al., 2009); (2) the inverse Logit (IL) method (Lindquist and Wager, 2005); (3) canonical correlation analysis (CCA) using a single-gamma function and its derivatives (Hossein-Zadeh et al., 2003a; Shams et al., 2006); (4) the basis expansion method using Gamma basis functions (BEG) (Prokopiou et al., 2019), and (5) the basis expansion method with spherical Laguerre functions (BEL) (Prokopiou et al., 2022, 2020). We further introduce a sixth method, an optimized CCA approach, which leverages the simplicity of the CCA method but allows for more flexibility in the basis sets. We directly compare these six methods under different ground-truth dCVR assumptions and different SNR conditions to identify their suitability for estimating different aspects of dCVR.

## Methods

### Evaluated modeling approaches

The deconvolution strategy assumes a linear time-invariant transfer (impulse response) function *h*(*t*) that relates the BOLD signal (*Y*(*t*)) and the CO_2_ fluctuations (*X*(*t*)).

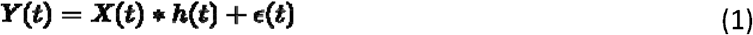

Where *h*(*t*) denotes dCVR in the remainder of this work, * denotes convolution, and *ε*(*t*) represents the residuals.

### Maximum a posteriori likelihood (MAP) method

This method has been extensively used to estimate physiological response functions in restingstate fMRI data data (Chang et al. 2009; Golestani et al. 2015). It has minimal model assumptions, with the exception of *h*(*t*) exhibiting a smooth characteristic in accordance with a Gaussian prior. Thus, we expect the performance of the MAP method to be most favourable in the case of the Gaussian ground-truth *h*(*t*). The response function *h*(*t*) can be solved based on Bayes’ Rule by maximizing *p*(*h*(*t*)|*Y*(*t*)),

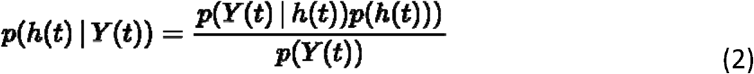

Here, *h*(*t*) is assumed to be a Gaussian process. That is,

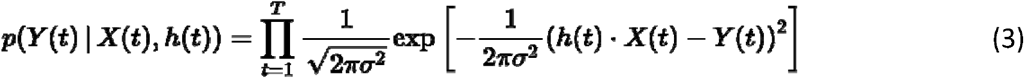

Thus,

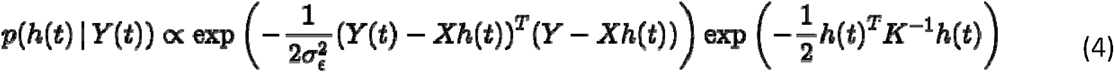

where *K* is the covariance matrix defined as

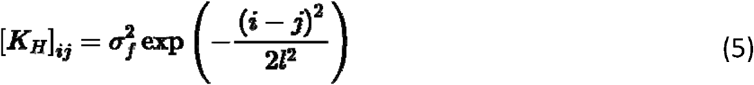

where *l* regulates the smoothness of *h*(*t*) and *σ_f_* regulates the distance between *h*(*t*) and its mean. The solution to *h*(*t*) becomes

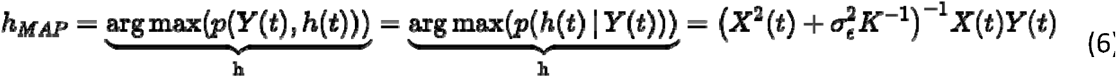

where *σ_ε_* is the variance of the BOLD time series *h*(*t*).

### Inverse Logit (IL) method

This method was applied as was described by Lindquist et al. (Lindquist et al., 2009), and will not be described at length here. The hemodynamic response is modeled as a linear combination of scaled and shifted inverse logit (IL) functions (Lindquist and Wager, 2007),

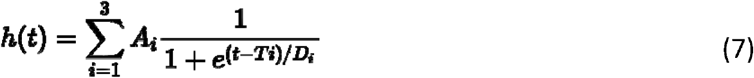

where *A_i_, D_i_, T_i_*, are constants to be determined. *A_i_*, controls the direction and amplitude of the curves, whereas *D_i_*, and *T_i_*, control the shift of their center and the angle of their slope, respectively. *A_i_*, is constrained such that *h*(*t*) starts and finishes at zero. The first IL function (Eqn. 7, *i* = 1) describes the rise of the positive lobe of *h*(*t*), the second IL function (*i* = 2) describes the fall, while the third IL function (*i* = 3) describes the appearance of the undershoot. Equating *A_1_* to *A_2_* and *T_1_* to *T_2_* results in a total of 7 unknown parameters (Shan et al., 2014). These parameters are estimated through a stochastic process such as simulated annealing, in which the best fit parameter set is found through random movement through the parameter space (Lindquist and Wager, 2007).

### Basis-expansion methods

In this work, we use a first-order Volterra kernel, which is equivalent to the impulse response. This has been found to well characterize dCVR (Prokopiou et al., 2019) instead of higher-order kernels, and can be estimated based on expansion of a basis set is chosen. The modeling problem becomes estimating the coefficients of the basis expansion *c_j1_* for the *1*^th^ basis function *j=1* for the impulse response) using ordinary least squares, where *1=0...L-1* (Marmarelis, 1993). In this work, two variants of the basis-expansion method are included, each using a different basis set.

### Basis expansion method with spherical Laguerre basis (BEL)

The spherical Laguerre is well suited to modeling an impulse response *h*(*t*) that starts from 0 and decay exponentially (Prokopiou et al., 2022). The Laguerre basis set has been used extensively to model physiological systems (Dabir et al., 2009; Francis et al., n.d.). The spherical Laguerre basis set is given by (Marmarelis, 1993; Prokopiou et al., 2022)(Leistedt and McEwen, 2012)

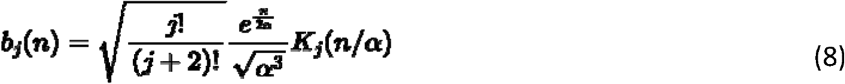

Where *j* = 0... *L-1*, and α is set to {2, 4} and to {0.5,1}. Furthermore,

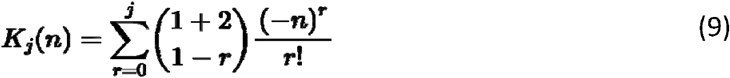

### Basis expansion method with Gamma basis (BEG)

As a reference to the spherical Laguerre approach, a Gamma basis was also used, defined by

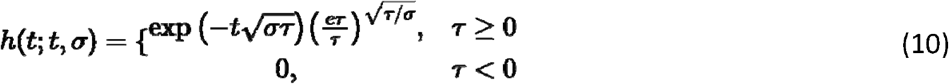

where *σ* represents the peak width (dispersion) and *τ* its location. The BEG method adopts the principle-components (PCA) basis approach (Aguirre et al., 1998; Woolrich et al., 2004), whereby an orthonormal basis set of the extended set of Gamma basis functions are determined from the top 2 singular values of the PCA, and used for modeling *h*(*t*), as described in (Prokopiou et al., 2019). For generating the full gamma basis set as input to the PCA, the range of σ and τ was [0.02 0.4] and [114], respectively.

### Canonical-correlation analysis (CCA) method

Canonical correlation analysis (CCA) is a simple and frequently used algorithm for finding the best linear combinations of two sets of multidimensional variables such that the correlation between the resultant basis vectors is mutually maximized. For details on its formulation, see Supplementary Materials.

### Optimized CCA method

If the *h*(*t*) subspace vectors are normalized prior to applying CCA, rescaling the canonical vectors has no effect on the resultant correlation for modeling purposes. Thus, we can rewrite the above resulting combination vectors that yield the highest correlation in terms of scaled vectors (*U*’ and *V*’):

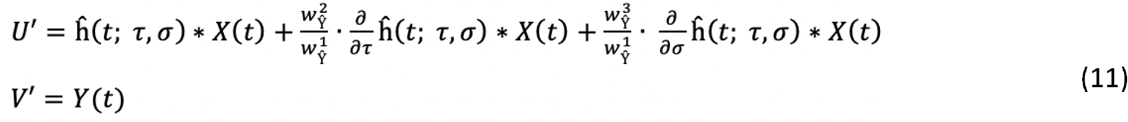

So we can rewrite Eqn. A3 as follows,

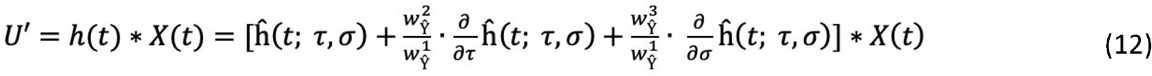

Since the above signal has the highest correlation with the BOLD signal, *h*(*t*) is the estimated dCVR. Through its ability to find the weights that maximize the correlation between two sets of signals, one of which was the BOLD signal and the other being the modeled signal, the CCA method also lends itself to impulse-response estimation.

Since conventional CCA uses the derivatives with respect to τ and σ, it can optimally find *h*(*t*) based on *ĥ*(*t*) as long as the parameters of the actual *h*(*t*) are close to the predefined parameters for *ĥ*(*t*). To address this fundamental limitation in the conventional CCA approach while taking advantage of its simplicity, we further propose an iterative optimization component to CCA based on Euler’s discretization method. The proposed method is based on the concept of derivatives, which implies that the small changes in a parameter of a function/model can be implemented with the linear combination of the model and the first derivative of the model with respect to the parameter. Here, for simplicity, we retain the single-gamma *h*(*t*) used in the initial CCA work (Hossein-Zadeh et al., 2003b), because of its simple implementation due to the small number of free parameters as well as the ability to cover a wide range of changes in the characteristics of *h*(*t*) through its use of derivative terms. Once the derivative of the initial *ĥ*(*t; τ,σ*) is determined with respect to parameters σ and τ, we can calculate *ĥ*(*t; τ,σ* + Δ*σ*) by multiplying the derivative, i.e., 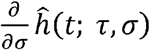, with a predefined or an adaptive step size, i.e., Δσ. This can be implemented by using Euler’s discretization:

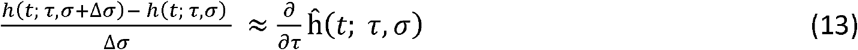

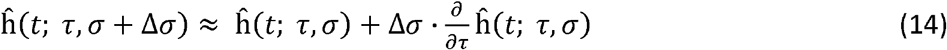

Accounting for the independence of the parameters (i.e. τ and σ), we can rewrite the above equation for two free parameters as follows:

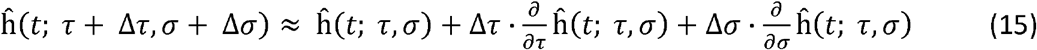

According to Euler’s theorem, the above formula is only asymptotically valid for small changes in each parameter, so the change step sizes (Δ*τ* and Δ*σ*) should be kept small enough 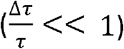. By keeping on repeating the above formula within a valid interval and direction for each parameter, we continue to update ĥ(*t*), and would reach a specific value for each of them so that the parameters of ĥ(*t*) are the best fit for the data.

Comparing Eqn. (A3) and (12), we can observe that their forms are nearly identical, except for the use of scaled coefficients (i.e., 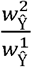 and 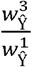) to determine the goodness of fit. We propose an adaptive optimization for CCA that iterates depending on the direction of the residual error. Positive values for these scaled coefficients indicate the need to increase *τ* and *σ* in *ĥ*(*t*) and negative values indicate the opposite. If these scaled coefficients are close to zero, it is suggested that ĥ(*t*) is a sufficiently accurate proximation of *h*(*t*), and the iteration can stop. However, due to the noise or lack of high correlation between the fMRI time series and the stimulus signal (e.g., PETCO_2_), changing the parameters in the direction and size obtained from the CCA may no longer increase the correlation, at which point the iterations would terminate. Moreover, for low SNRs, the stopping point can also be a local instead of global minimum. The robustness of this approach for dCVR modeling has yet to be determined.

Since correlation is scale insensitive, we can first prioritize fitting for the shape of *h*(*t*), and then estimate its amplitude by comparing *Y*(*t*) with the convolution of *X*(*t*) with *h*(*t*). Next, to further minimize the effect of noise, we considered only the data points of *Y*(*t*) and *X*(*t*) that are in their respective top quartiles in terms of magnitude. Then, we convolve *h*(*t*) and *X*(*t*), and divide the result by *Y*(*t*) to produce a ratio time series. The average of this ratio time series is multiplied to *h*(*t*) to scale its amplitude to match that of *Y*(*t*).

### Simulation-based validation

We used simulated data sets to evaluate the performance of all methods in terms of their abilities to accurately and robustly estimate multiple characteristics of dCVR under a broad range of SNR conditions.

### Ground-truth CO_2_ response functions

We employed 4 ground truth dCVR functions selected from the literature: 1) single Gamma, 2) double Gamma, 3) triple inverse logit, and 4) Gaussian, with a specific range of timings and amplitudes. These ground-truth dCVRs are plotted in Fig. 1. All dCVR ground-truth forms are assumed to exhibit zero arrival delay. 40 variations are chosen for each of the four forms, with representative examples plotted in Fig. 1b-d. As the shape of dCVR can vary widely between health and disease, the dCVR timings are chosen to accommodate a wide range of physiologically plausible shapes reported in previous literature, including those based on neuronal activation (Glover, 1999; Lu et al., 2007; Prokopiou et al., 2022)) and CO_2_ (Golestani et al., 2015; Prokopiou et al., 2019). Moreover, the ground-truth *h*(*t*) are not chosen to favour any parameter ranges assumed in the methods tested.

1. The single Gamma *h*(*t*) was modeled according to the following Gamma function (Lange and Zeger, 1997, Hossei-Zadeh et.al., 2003 and Friston et. al. 1998):

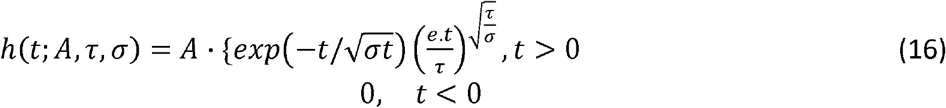

where A controls the height (maximum amplitude) of *h*(*t*) and τ and σ control its latency (time to peak) and shape (mostly the width), respectively. To model the *h*(*t*) variations, we randomly selected τ and σ in the ranges of [3, 9] and [0.05, 0.5], respectively.
2. The double Gamma *h*(*t*) was modelled by the sum of two gamma functions as follows (Shan et al., 2014).

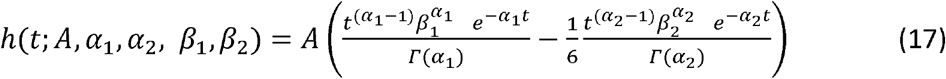

where *Γ* is the gamma function. *α*_1_ and *α*_2_ respectively specify the peak times of the first and second Gamma functions while *β*_1_ and *β*_2_ control the shape of the first and second gamma functions. We randomly selected *α*_1_ *α*_2_ *β*_1_ and *β*_2_ in the range of [3, 9], [6, 25], [0.5, 2], and [0,1.5], respectively (close to the ranges specified in Shan et. al 2014).
3. The inverse logit *h*(*t*) was modelled by the following formula

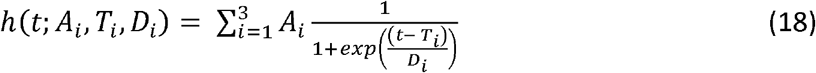

where *T_i_* and *D_i_* control the shift center and slope of each of the logit functions. We randomly chose *D_i_*, *T*_1_, *T*_2_, and *T*_3_ in the range of [1, 5], [1, 7], [2.5, 17.5], and [3.25, 22.75], respectively, such that the general form of *h*(*t*) is maintained. We also randomly selected *A*_1_ and calculated *A*_2_ and *A*_3_ based on the formulation proposed in (Lindquist et. al., HBM, 2007).
4. The Gaussian *h*(*t*) was modeled by the well-known formula of the Gaussian distribution

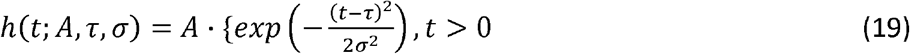

where *A* = 1, τ and σ control the delay (time to peak) and the FWHM of respectively. We randomly selected the τ and σ to be in the ranges of [5,12] and [1, 6], respectively.

**Figure 1.**
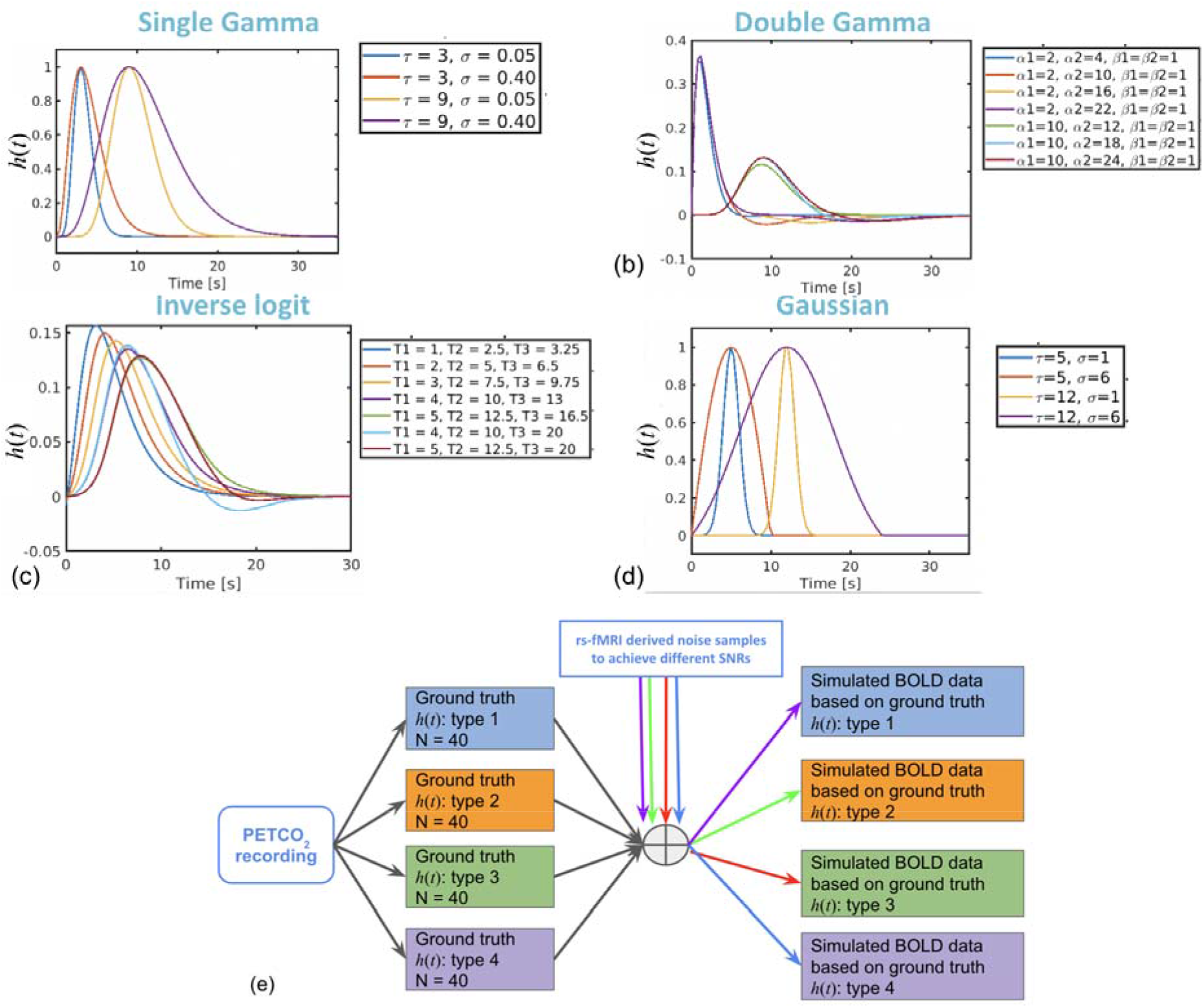
Summary of simulation framework. A total of 4 commonly assumed groundtruth shapes are used in turn (a-d) to convolve the end-tidal CO_2_ time course, and noise is added to the result at designated SNRs to produce the test data (e).

### Noise levels

We prepared different sets of simulated data using different ground truth *h*(*t*) and SNR values. The procedure is illustrated in Fig. 3. We added noise with scaled amplitude to the results of the convolution, aiming to acquire voxel-wise SNRs of −10, −6.9, −3.0, 0, 3.0, and 6.9 dB respectively.

**Figure 3.**
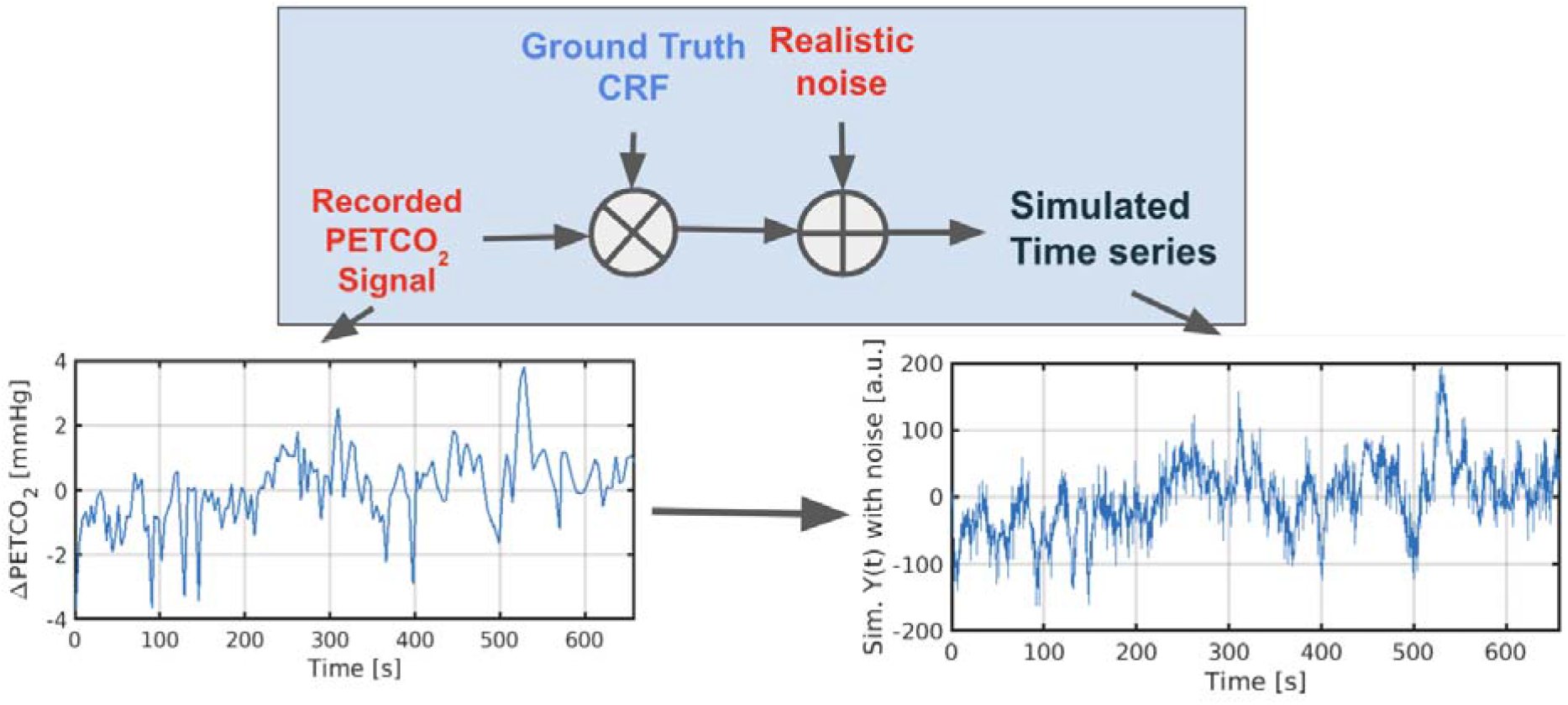
Addition of noise to achieve different SNRs. A sample PETCO_2_ and resultant noise-added BOLD time course are shown.

Here, SNR is defined as 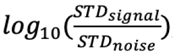, where STD represents the standard deviation. As the signal contribution is a waveform computed by convolving resting PETCO2 with a known, SNR is computed using signal STD instead of amplitude. To maximize the similarity between the generated signal and the signals acquired from the fMRI scanner, we first divided the experimental data set into two groups (Groups 1 and 2). Then, we used BOLD signals from the 340 voxels located in the gray matter of the Group 1 data set to generate realistic fMRI noise, while the PETCO2 signal used for that set was randomly selected from Group 2 and then permuted.

To generate realistic noise time series for our simulations, we randomly sampled greymatter voxels from resting-state fMRI data acquired from a healthy young volunteer. All data were acquired on a Siemens TIM Trio 3 Tesla System (Siemens, Erlangen, Germany), with 32-channel phased-array head coil reception and body-coil transmission. During the scans, participants were instructed to be relaxed and keep their eyes closed. Slice-accelerated single shot gradient-echo (GE-EPI) images (Feinberg et al., 2011; Setsompop et al., 2011) were acquired (TR = 389 ms, TE = 30 ms, flip angle = 40°, 15 slices, 3.44 × 3.44 × 6 mm3, 2230 volumes, acceleration factor = 3, phase encoding shift factor = 2) performed in an inter-leaved fashion. For all participants, we also collected T1-weighted anatomical images for anatomical registration and segmentation (MPRAGE, TR = 2400 ms, TE = 2.43 ms, FOV= 256 mm, voxel size = 1 × 1 × 1 mm3) in the same session. For physiological monitoring, we recorded PETCO_2_ signals using a RespirAct™ system (Thornhill Research Inc, Toronto, Canada).

All calculations were performed using in-house scripts written in Matlab 2019 (Mathworks, Natick, Massachusetts, USA).

### Evaluation criteria

#### dCVR parameters

To facilitate a comprehensive assessment, each estimated *h*(*t*) is characterized by the following 5 parameters:

1. The time to peak (TTP): the time it takes for *h*(*t*) to reach its positive peak;
2. The time to half maximum (TTH): the time for *h*(*t*) to decrease to the half of its peak;
3. The full-width-at-half-maximum (FWHM): the width of *h*(*t*) measured at the points of half-maximum, used to describe dispersion effects in dCVR.
4. The area under *h*(*t*): corresponds to the steady-state BOLD signal response to an impulse PETCO_2_ input, and is thus defined as the static CVR;
5. The correlation coefficient between the ground-truth and estimated *h*(*t*): this assesses the agreement in overall shape of *h*(*t*).

#### Estimation accuracy and variability

For each assumed ground-truth *h*(*t*) and at each SNR, these 5 metrics were assessed and the results were summarized through the following performance metrics:

- Relative fractional error:

◦ estimated/ground-truth x 100%
- Absolute fractional error:

◦ |estimated |/ground-truth x 100%
- Computational time:

◦ The time taken to estimate a single *h*(*t*) averaged across all SNRs and ground truths; this is assessed on a 1.4 GHz Quad-Core Intel CPU with 2 GHz 8GB memory.

Furthermore, to further consolidate these diverse quality metrics, we defined a composite quality metric as illustrated in **Fig. 5.** This composite metric not only depends on the mean estimation accuracy (Accuracy Index) but also depends on the variability of the estimation accuracy across SNRs and ground truths (Robustness Index). That is, a high composite quality metric indicates that a method not only produces the highest average estimation accuracy in terms of all parameters, but is also the least variable in terms of performance.

**Figure 4.**
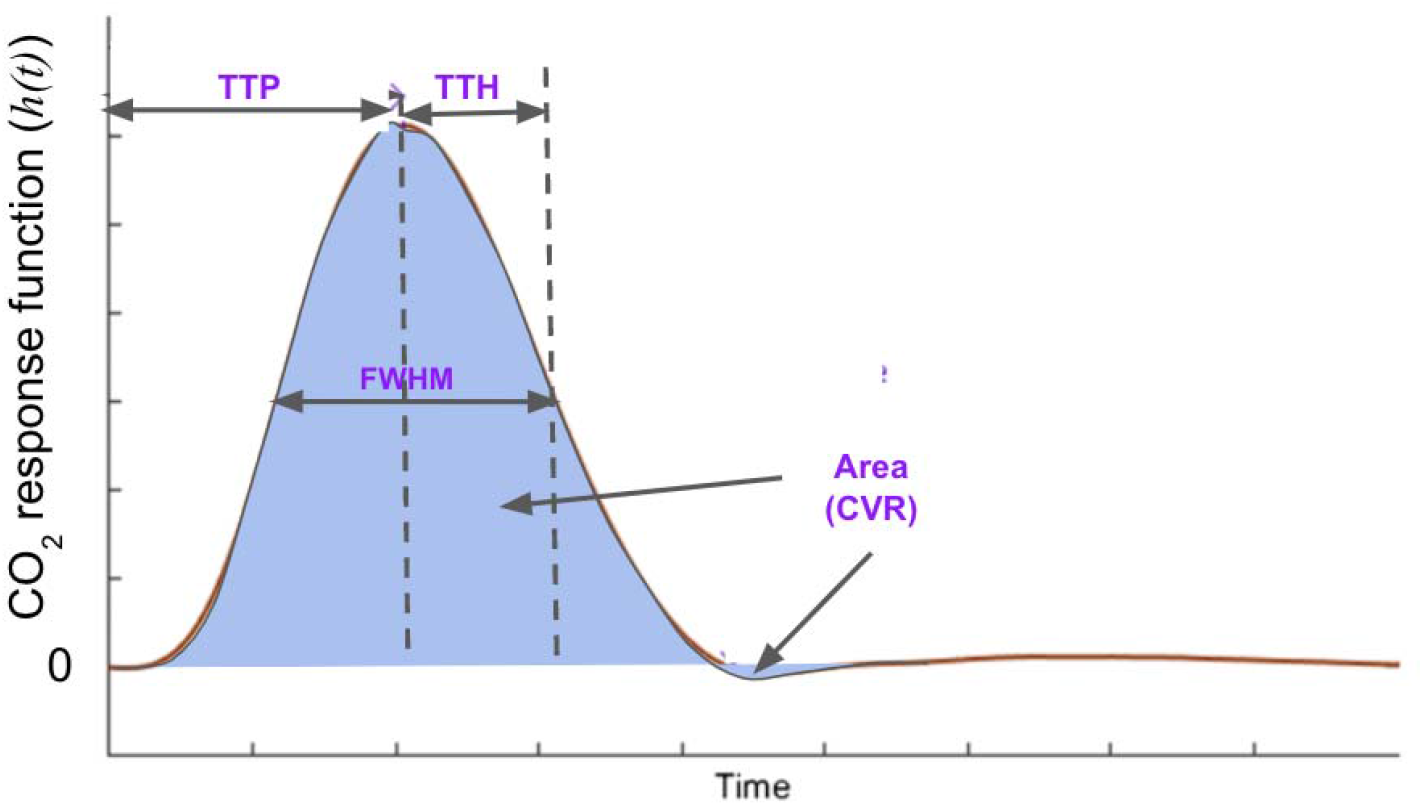
Illustration of the dCVR metrics. TTP: time-to-peak; TTH: time-to-half-max; FWHM: full-width at half-maximum; area: the area of *h*(*t*), equivalent to the CVR amplitude.

**Figure 5.**
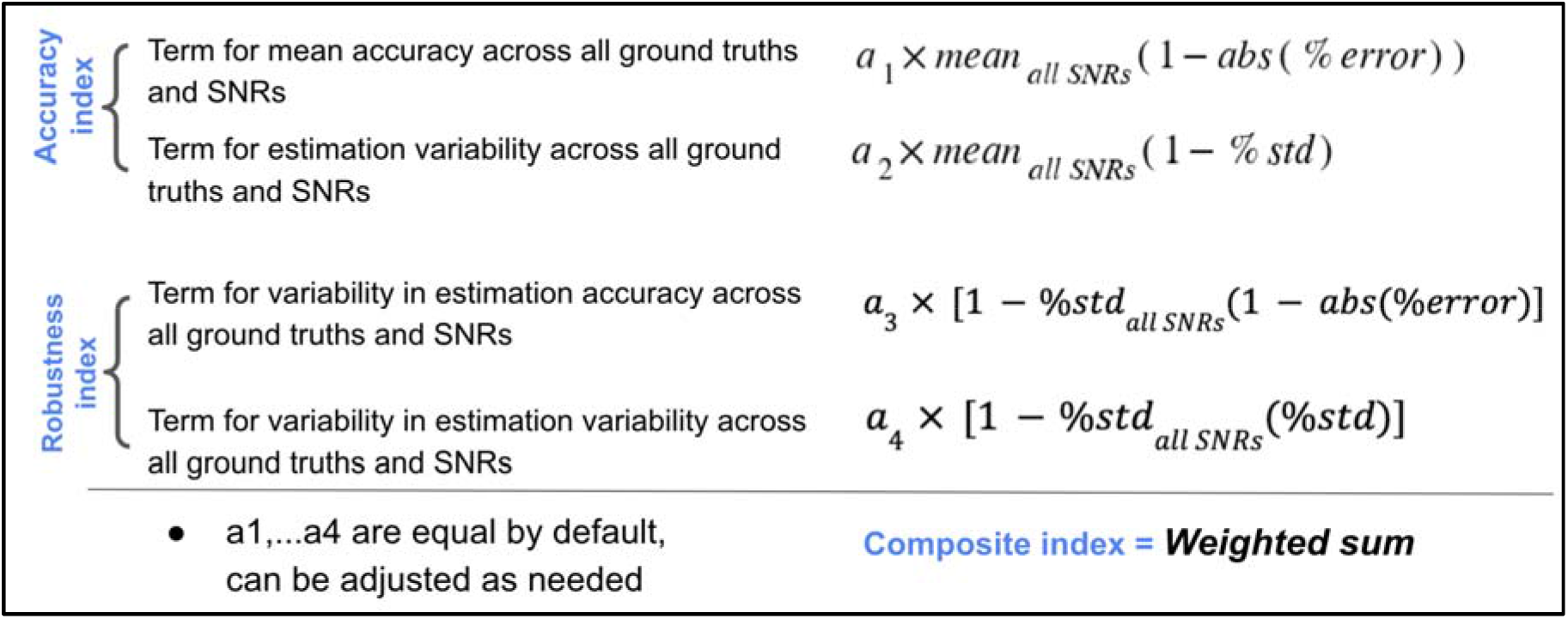
Illustration of the composite quality metrics. The Accuracy Index contains a penalty term for low *h*(*t*) estimation accuracy, whereas the Ro bustness Index is penalized for high *h*(*t*) estimation variability (in spite of high average accuracy). The composite index embodies both aspects.

## Results

As can be seen in **Fig. 6,** TTP estimation accuracy of different methods exhibits limited dependence on the assumed ground truth. For instance, the CCA method generated the highest TTP estimation error for all ground truths, with the exception of the double-Gamma ground truth, where the MAP method generated the highest TTP error (Fig. 6b). On the other hand, TTP estimation error decreases with increasing SNR for all methods except the CCA method. Furthermore, the estimation errors are distributed about zero (Fig. 6e-g) except for the Gaussian ground truth, where all methods systematically overestimate TTP (positive relative error). The methods generating the lowest TTP estimation errors are the IL, the optimized CCA (CCA_opt_) and the BEL methods.

**Figure 6.**
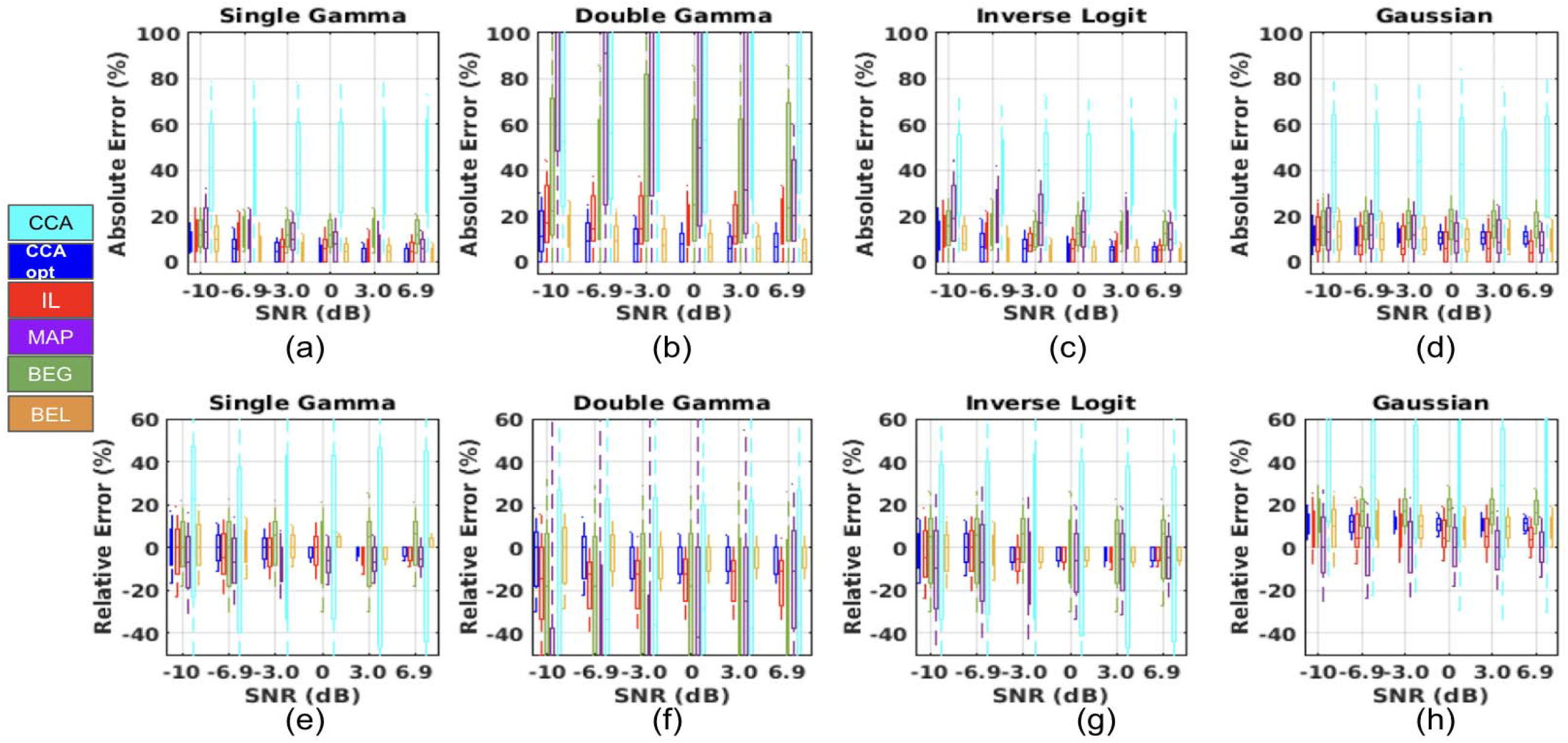
TTP estimation accuracy across all SNRs and ground truth *h*(*t*) forms. The absolute (a-d) and relative errors (e-h) in TTP estimation for each method is plotted as box plots across different SNR values, whereby the first to third quartile of TTP estimation error across the 300 simulated BOLD signals is shown by the extent of the boxes.

As shown in **Fig. 7,** like TTP, TTH estimation accuracy also exhibits limited dependence on the assumed ground truth. That is, with the exception of the double-Gamma ground truth, where the MAP method generated the highest TTH error (Fig. 7b), the CCA method performs worst irrespective of ground truth *h*(*t*). Moreover, similar to TTP, TTH estimation error also decreases with increasing SNR for all methods except the CCA method. Unlike TTP, TTH is generally overestimated by the BEL method (positive relative error in Fig. 7e, g, h) and underestimated by the CCA method (negative relative error in Fig. 7e-h). Once again, the methods generating the lowest TTH estimation errors are the IL, the optimized CCA (CCA_opt_) and the BEL methods.

**Figure 7.**
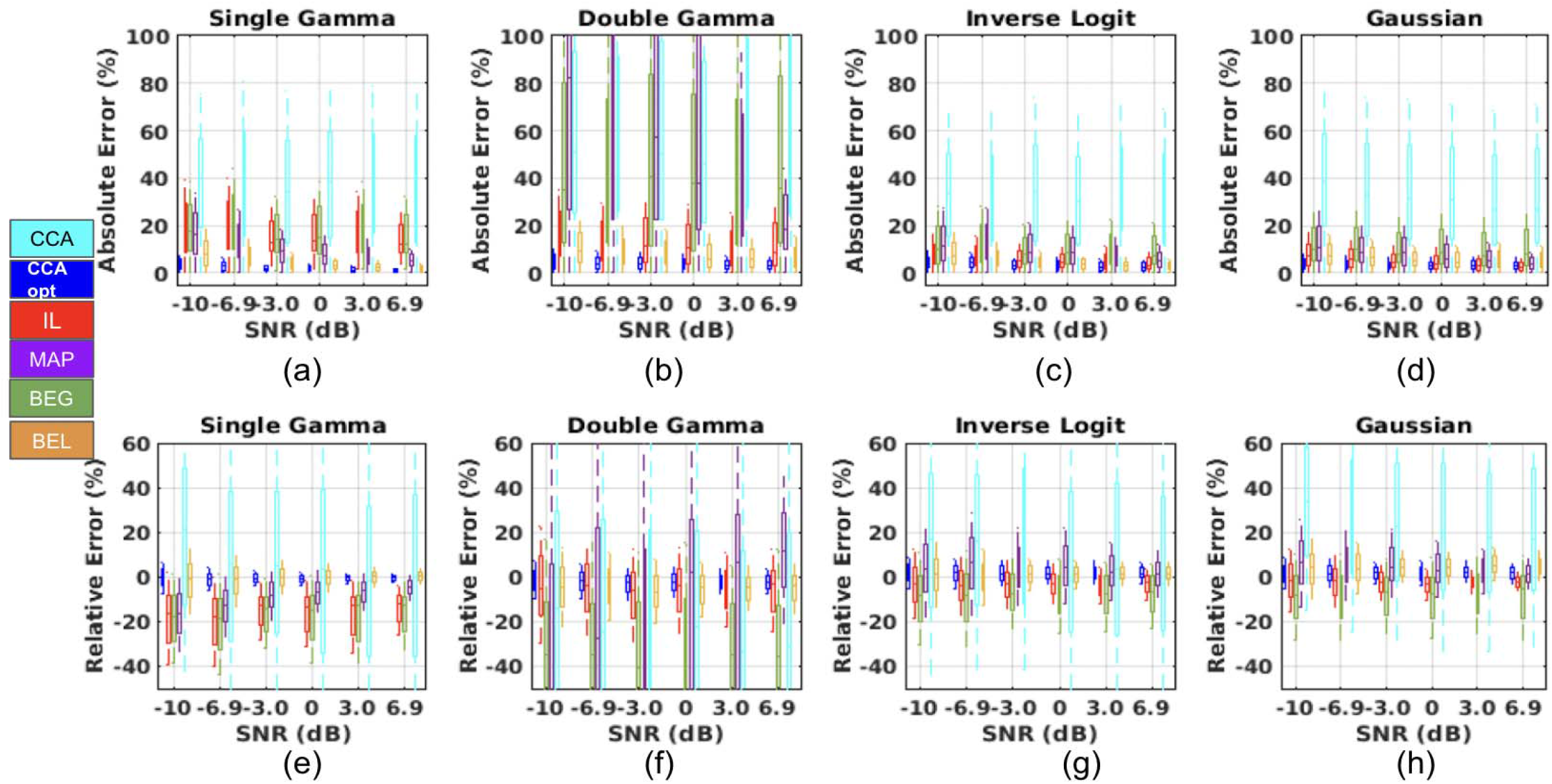
TTH estimation accuracy across all SNRs and ground truth forms of *h*(*t*). The absolute and relative errors in TTH estimation for each method is plotted as box plots across different SNR values, whereby the first to third quartile of TTH estimation erro r across the 300 simulated BOLD signals is shown by the extent of the boxes.

As shown in **Fig. 8,** compared to TTP and TTH, FWHM estimation accuracy exhibits stronger dependence on the assumed ground truth *h*(*t*). For instance, the IL method heavily underestimates the FWHM but only for the single-Gamma ground truth (Fig. 8e), while the BEG method heavily over-estimates the FWHM but only for the double-Gamma ground truth (Fig. 8f). The CCA method is no longer the worst performer - replaced by the BEG method (with the highest estimation error, Fig. 8a & b). Moreover, FWHM estimation error generally decreases with increasing SNR for all methods except the CCA and BEG methods. The methods associated with the lowest FWHM estimation errors across all ground truths and SNRs are the BEL and CCA_opt_ methods.

**Figure 8.**
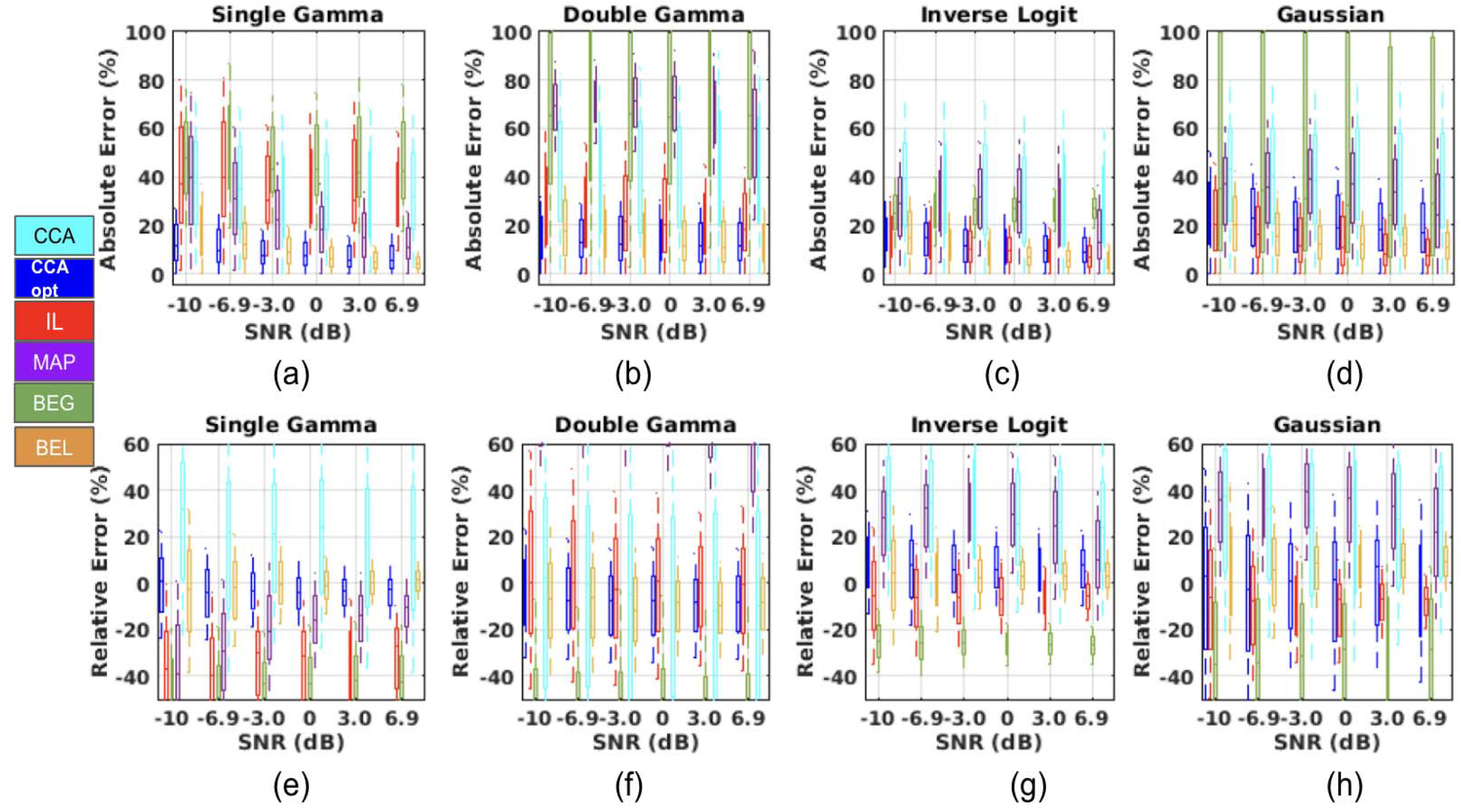
FWHM estimation accuracy across all SNRs and ground truth forms of *h*(*t*). The absolute and relative errors in FWHM estimation for each method is plotted as box plots across different SNR values, whereby the first to third quartile of FWHM estimation error across the 300 simulated BOLD signals is shown by the extent of the boxes.

The area of the estimated *h*(*t*) (equivalent to the CVR amplitude) is the only shape-independent parameter of *h*(*t*) assessed in this work. As shown in **Fig. 9,** CVR estimation variability decreases with increasing SNR for most methods (Fig. 9a-d). Moreover, the CVR amplitude is best estimated using the IL and BEL methods (Fig. 9a-d), followed closely by the CCA_opt_ method. Of these, the IL method yielded the lowest variability, sustained over all SNRs and ground-truth dCVRs. The MAP and CCA (errors too large to be shown) methods yielded the largest errors. While each model was expected to perform best for its respective assumed *h*(*t*) shape, this was not the case, as the variability of all methods was highest for the double-Gamma ground truth (Fig. 9b).

**Figure 9.**
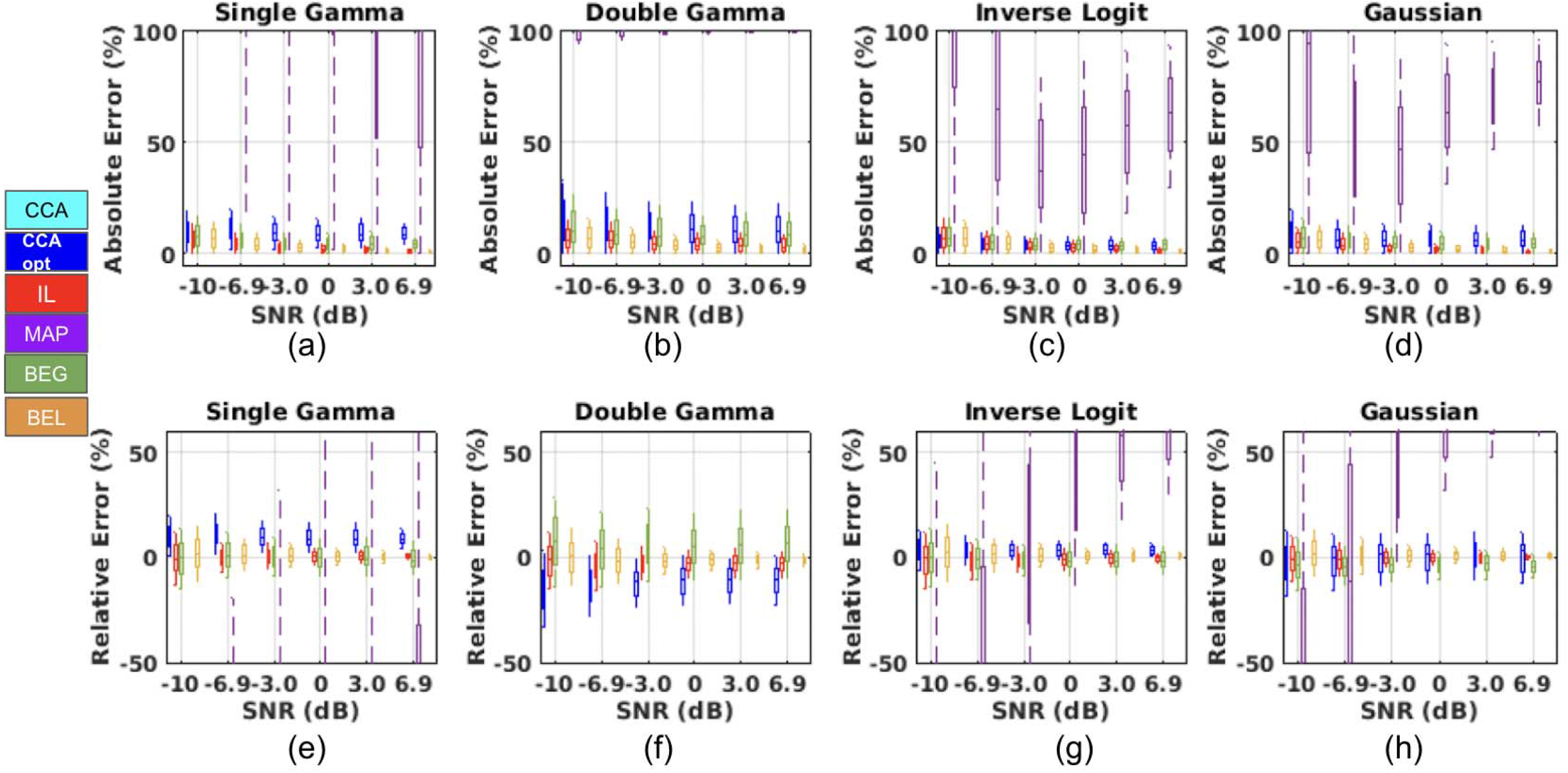
CVR (area of *h*(*t*)) estimation accuracy across all SNRs and ground truth forms of *h*(*t*). The absolute and relative errors in CVR estimation for each method are plotted as box plots across different SNR values, whereby the first to third quartile of CVR estimation error across the 300 simulated BOLD signals is shown by the extent of the boxes. The absolute errors associated with the CCA method are too large to be shown.

As shown in **Fig. 10,** considering a correlation coefficient of 1 to represent a perfect shape match between the estimated and true *h*(*t*), the shape of *h*(*t*) is best estimated by the IL, BEL, BEG and CCA_opt_ methods, and least well by the CCA and MAP methods. Of these, the CCA_opt_ and IL method yield the lowest variability, and this is sustained over all SNRs and ground truth forms of *h*(*t*). The correlation with ground truth approaches 1 with increasing SNR. The performance of different method in terms of correlation appears not to depend heavily on the assumed ground truth. As a case in point, the MAP method, which assumes a Gaussian ground truth, yielded similar error levels for the Gaussian and IL *h*(*t*) (Fig. 10 c, d). Here again, the variability of all methods was highest for the double-Gamma ground truth (Fig. 10b), instead of being biased by their respective model assumptions.

**Figure 10.**
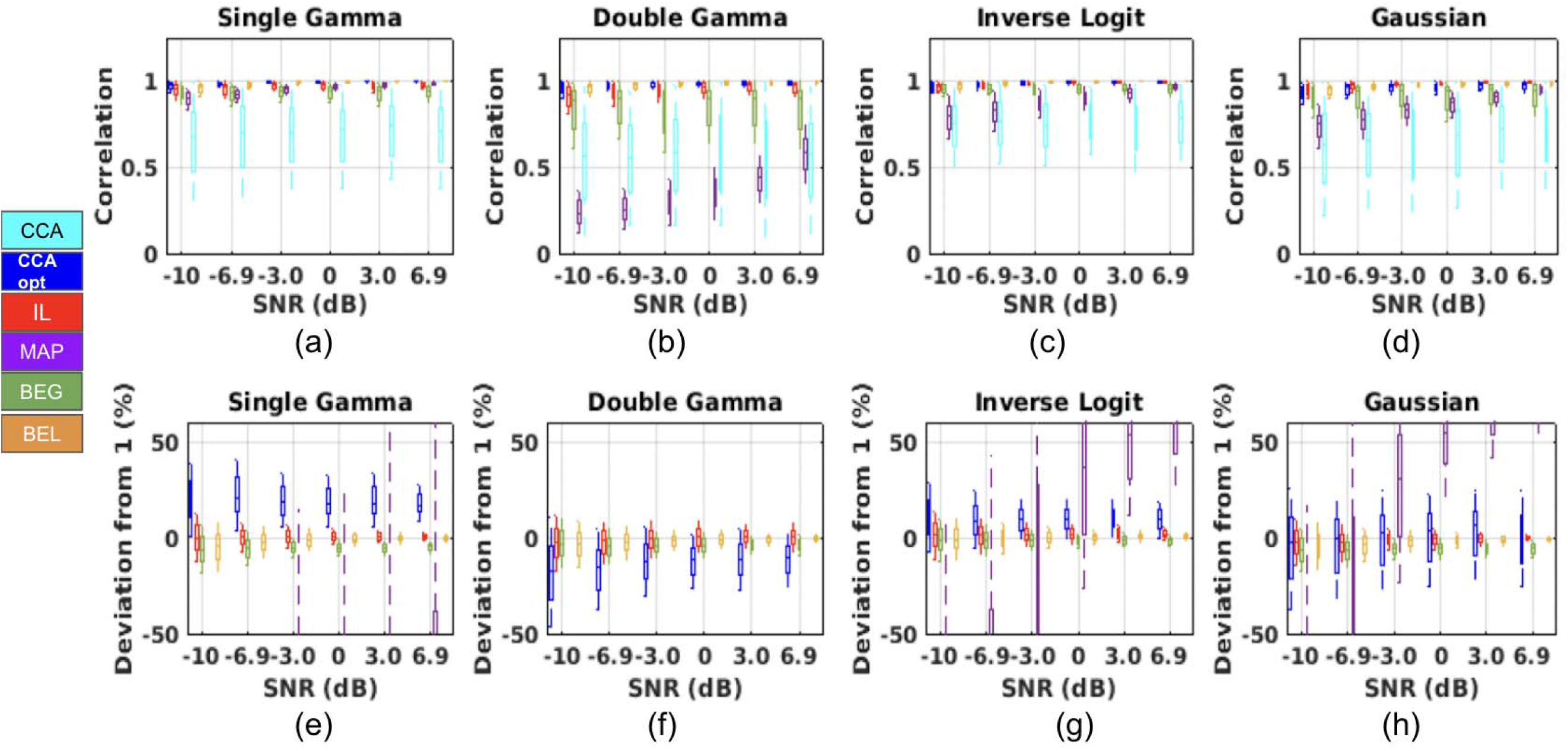
Correlation coefficients between the estimated and true ground truth dCVRs across all SNRs and ground truth forms of *h*(*t*). The correlation coefficients (a-d) and their relative deviations from the ideal correlation coefficient of ‘1’ (e-h) for each method is plotted as box plots across different SNR values, whereby the first to third quartile of the correlation (and corresponding relative errors) across the 300 simulated BOLD signals is shown by the extent of the boxes. In (e)-(h), The absolute errors associated with the CCA method are too large to be shown.

As shown in **Fig. 11,** different methods are associated with very different Accuracy and Robustness Indices **(Fig. 11a & b).** The BEL method is associated with the highest accuracy across all dCVR (h(t)) metrics, while the MAP and CCA methods are associated with the worst accuracy (Fig. 11a). In terms of robustness, MAP is associated with the worst performance **(Fig. 11b).** Combining accuracy and robustness, the IL, BEL and CCA_opt_ methods perform similarly **(Fig. 11c).** Overall, the results suggest that different dCVR metrics are best estimated using different methods. For instance, the IL method produces worse FWHM estimates than the CCA_opt_ and BEL methods, while the BEL method produces better TTP and CVR estimates than the others.

**Figure 11.**
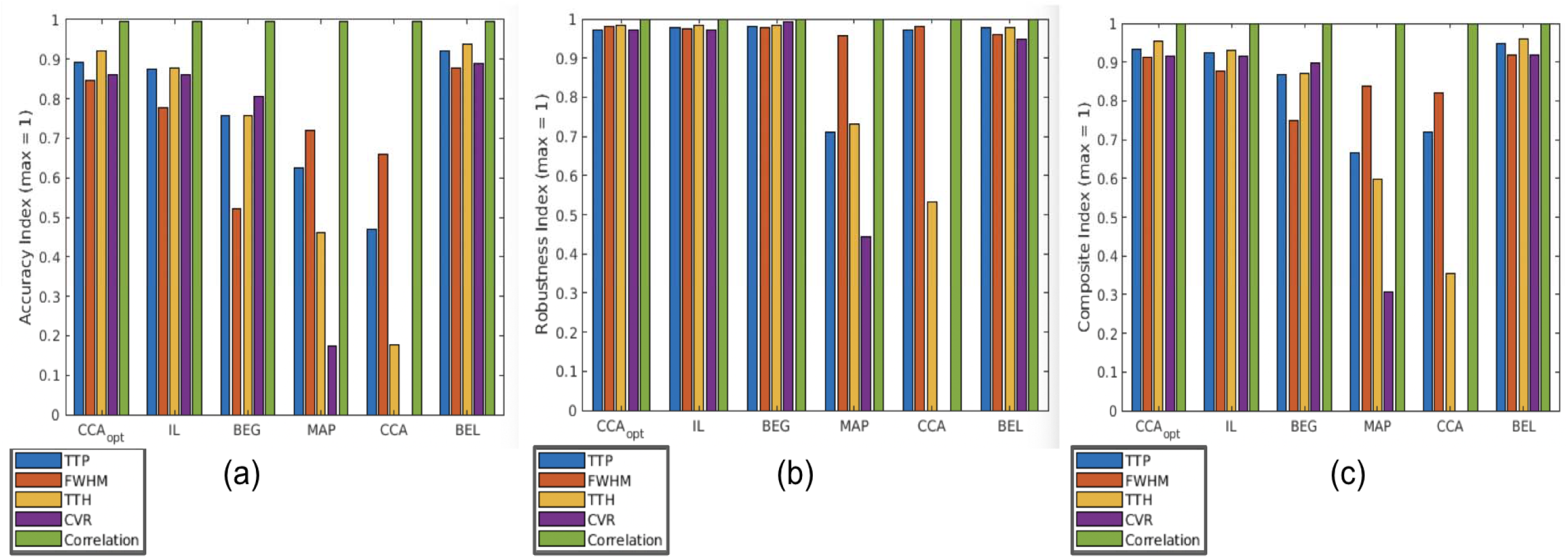
Summary of composite quality metrics, shown for different dCVR (*h*(*t*)) metrics, including TTP, TTH, FWHM and area (CVR), and correlation, for all methods.

Lastly, in terms of computational complexity, the processor times for all methods are summarized in Table 2. The CCA-based methods are by far the fastest, taking 0.0011 s and 0.0013 s per execution for CCA and CCA_opt_, respectively. They are followed by the BEG and MAP methods, which range between 0.0027 and 0.0058 s. The IL method is the next longest, taking 1.35 s per execution, and finally, the BEL method takes 3.85 s per execution.

**Table 1.**
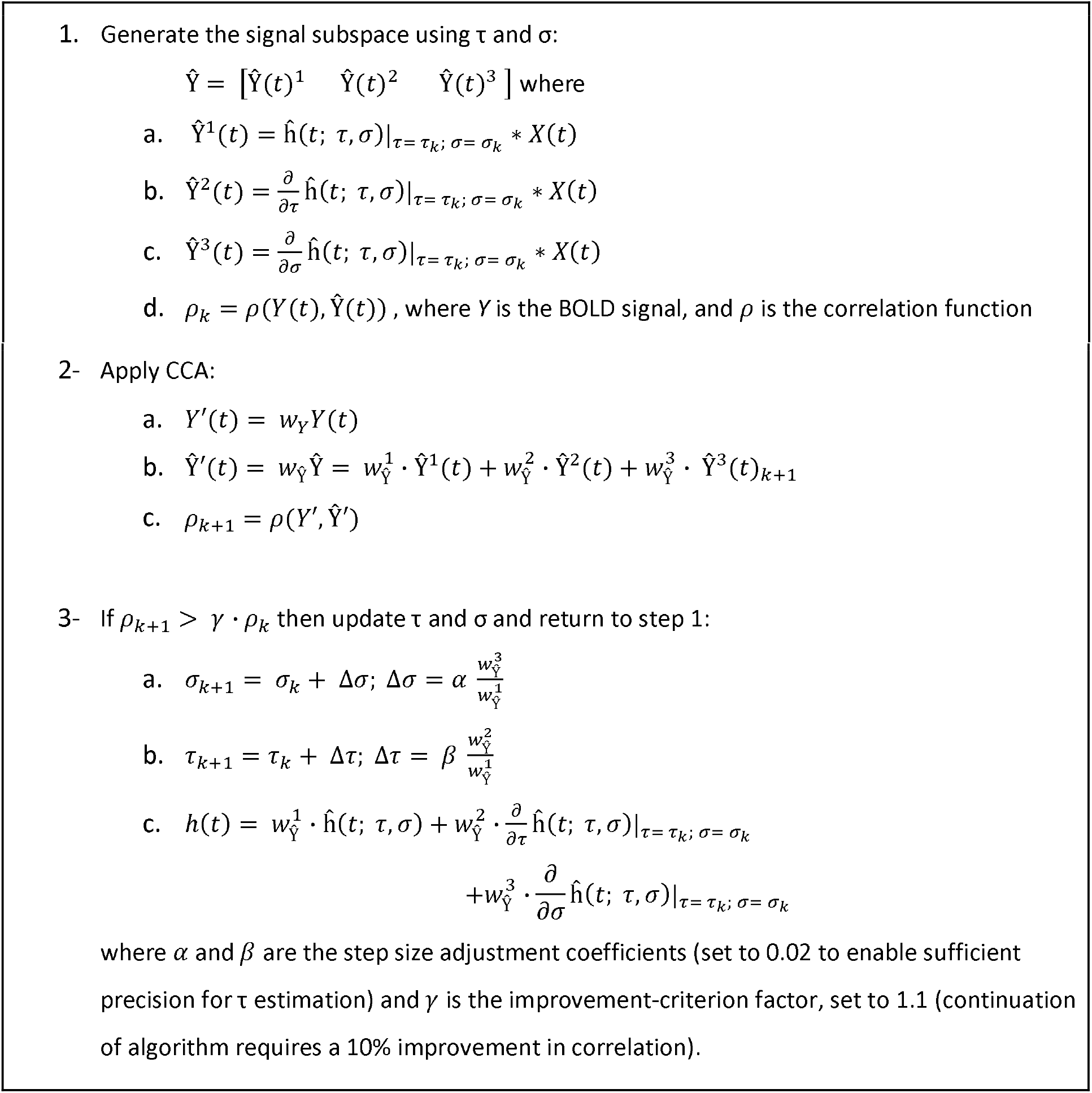
Summary of the proposed adaptive optimized CCA (CCA_opt_) algorithm. The optimization step (step 3) distinguishes the CCA_opt_ method. At each iteration round, CCA is applied to the signal subspace generated based on the present parameters (τ and σ), and then the parameters evolve based on CCA coefficients.

**Table 2.** Summary of computation time for all methods. Timing values are specified in seconds for each time course, averaged over all SNRs and ground-truth scenarios. The normalized average (norm. average) allows for a generalizable processing-time assessme nt, as processors may differ widely. The reference (normalization factor) is the timing for the IL method.

| ***Mean±std (s)*** | **CCA<sub>opt</sub>** | **IL** | **BEG** | **MAP** | **CCA** | **BEL** |
| --- | --- | --- | --- | --- | --- | --- |
| Single Gamma | 0.0015±0.0012 | 1.43±0.48 | 0.0041±0.010 | 0.0058±0.0060 | 0.0011±0.0013 | 3.80±0.30 |
| Double Gamma | 0.0012±0.0002 | 1.33±0.26 | 0.0043±0.011 | 0.0038±0.0008 | 0.0011±0.0007 | 3.70±0.27 |
| IL | 0.0012±0.0006 | 1.37±0.21 | 0.0036±0.0008 | 0.0034±0.0004 | 0.0012±0.0010 | 3.85±0.38 |
| Gaussian | 0.0015±0.0008 | 1.27±0.36 | 0.0045±0.0007 | 0.0027±0.0004 | 0.0010±0.0002 | 2.06±0.60 |
| **Average (s)** | **0.0013±0.0007** | **1.35±0.30** | **0.0041±0.006** | **0.0039±0.0019** | **0.0065±0.0008** | **3.85±0.39** |
| **Norm. average** | **0.005** | **1.0** | **0.03** | **0.01** | **0.005** | **3.0** |

## Discussion

Modeling of dCVR can play a key role for understanding vascular physiology (West et al., 2019)(Holmes et al., 2020). However, due to the low signal-to-noise (SNR) conditions, especially in rs-fMRI, deconvolution of CO_2_ fluctuations from the fMRI signal is non-trivial. To overcome the uncertainties associated with the low SNR and the fact that the CO_2_ response can overlap in frequency with the neuronal hemodynamic response, model-based methods are generally more robust (Chen et al., 2005), but with the caveat that their performance may depend on the underlying model structure. In such cases, model simplicity and flexibility are competing aspects that dictate model accuracy, uncertainty and computational efficiency. In this work, we present a comparison of several recently published model-based deconvolution approaches for estimating *h*(*t*), including maximum a posterior likelihood (MAP), inverse logit (IL), canonical correlation analysis (CCA), and basis expansion (using Gamma and Laguerre basis sets). To this end, we devised a novel simulation framework that allowed us to target a wide range of SNRs, ranging from 10 to −7 dB, representative of both task and resting-state CO_2_ changes. In addition, we built ground-truth *h*(*t*) into our simulation framework, overcoming the practical limitation in methodological evaluation that the true *h*(*t*) is unknown. Moreover, to best represent realistic noise found in fMRI scans, we extracted it from in-vivo resting-state scans. Furthermore, we introduce a simple optimization of the CCA method (CCA_opt_) and compare its performance to these existing methods.

In this work, we demonstrate that dCVR can be extracted using modelling methods even from extremely noisy data. Moreover, the three leading methods are the IL (inverse logit) method, the BEL (basis expansion with spherical Laguerre functions) method and the CCA_opt_ (proposed optimized CCA) method. Notably, the CCA_opt_ and BEL methods were best at estimating dCVR timing parameters (TTP, TTH, FWHM), whereas the BEL and IL methods were best at estimating the CVR amplitude. The correlation between the estimated and true dCVR was highest for the BEL, IL and CCA_opt_ methods, irrespective of the assumptions for the true dCVR underlying each method. Lastly, of the three top-performing methods, the CCA_opt_ method, though not outperforming the BEL method in accuracy and robustness, requires less than 1/100^th^ of the computational time to execute. Thus, the choice of method may also depend on the dCVR parameter(s) of interest and the available computational resources, and each method has unique advantages and limitations.

### Features of dCVR

To characterize the dynamic attributes of dCVR, we did not adopt an exponential approach (Poublanc et al., 2015), as the time constant of the exponential can be very sensitive to the location of the noise floor, especially under low-SNR conditions. Instead, we defined timing parameters with respect to the peak of *h*(*t*), as was done in past literature (Li et al., 2019; Mayer et al., 2014; Rangaprakash et al., 2021).

The TTP represents the speed of dCVR on the rising edge. Although all ground-truth forms of *h*(*t*) are simulated with zero arrival latency, the TTP (time-to-peak) can also embody arterial arrival delay in realistic scenarios. In fact, the arterial delay can be sensitive to aging as well as diseases such as carotid stenosis and impaired collateral circulation (K. Chen et al., 2021; Ishii et al., 2020; West et al., 2019), where the CO_2_ response can start late with or without being sluggish. However, if a prolonged TTP is accompanied by a normal TTH, then the vascular response may be unaffected in spite of a pronounced arterial delay. As the TTH reflects the speed of dCVR on the falling edge, it also reflects the vasoconstrictive capacity, which can be pronounced in the presence of impaired vasodilation (Roustit et al., 2011). To complement TTP and TTH, the FWHM is taken to represent the duration of the dCVR, and also reflects, indirectly, our ability to detect the true peak of dCVR.

The area under *h*(*t*), in a linear response framework, i.e. the steady-state response to an external step CO_2_ stimulus, which is typically used as a quantitative measure of static CVR. Thus, our CVR value can in theory be either positive or negative, although negative CVR was not included in our simulations (as it is usually not expected, except in special cases - e.g. vascular steal (Mandell et al., 2008; Prokopiou et al., 2019; Sobczyk et al., 2014)). The clinical and research significance of quantitative CVR have been well established, and will not be repeated here (Blockley et al., 2017; Chen, 2018; Fierstra et al., 2013; Pillai and Mikulis, 2015; Pinto et al., 2020). However, the advantages of extracting dCVR shape parameters, and their potential improved sensitivity, are starting to be recognized in the study of aging and dementia (Gokcal et al., 2022; Holmes et al., 2020).

The correlation coefficient is a holistic shape metric. Unlike the other dCVR metrics, it is not interpretable physiologically in its own right. However, a high correlation reflects an accurately determined *h*(*t*) that reduces the occurrence of “false-positives” or “false-negatives” in terms of identifying the BOLD signal variations attributable to CO_2_ fluctuations.

### Performance of modeling methods

This study was motivated by the desire to model the CO_2_ response function under both highland low-SNR conditions, including those in resting-state fMRI. The challenge is magnified as the noise may overlap with signal, rendering it challenging to use filtering-based approaches such as the Wiener filter. In this regard, model-based methods have a unique advantage in that they can in theory extract the required response based on its shape, irrespective of spectral overlaps with signals of non-interest.

Though the MAP method is not model-independent, it is the most flexible of all approaches tested, as it is based on finite-impulse response filters and only assumes that the response has a Gaussian prior (Goutte et al., 2000). However, the flexibility of this approach may expectedly lead to underperformance under low-SNR conditions. While it is the most flexible of all methods tested and has been previously used for dCVR modeling in the resting state (Chang et al., 2009; Golestani et al., 2015), its performance is strongly SNR-dependent, as shown in **Figs. 6–10**. This suggests that the addition of certain model constraints can serve to moderate SNR dependence and improve a method’s noise immunity.

The IL method, in contrast, is highly parameterized and exhibited both remarkable accuracy and SNR insensitivity. The IL method was first introduced as a way to disentangle colinear model features (such as the peak response and undershoot in the canonical Gamma model) and maximize statistical power in general-linear model analysis of fMRI data (Lindquist and Wager, 2007). Moreover, the IL model parameters all have physiological interpretations, but the sheer number of fitted parameters also means the performance of the IL method is heavily dependent on an efficient fitting algorithm. In this work, we implemented the 7-parameter approach with model fitting based on the recommended simulated annealing approach (Lindquist et. al., HBM, 2007).

Both based on the basis-expansion framework, the BEG adopted Gamma basis functions while the BEL uses spherical Laguerre basis (Prokopiou et al., 2022, 2019). The performance of the two variations differed tremendously; the BEL method delivered the top performance of all methods tested, decisively leading the BEG method. This is likely due to the fact that in the former case, the extended gamma set is created empirically using prior information with regard to the underlying parameter space of the gamma basis set, which is held constant. In the work of Prokopiou et al. (Prokopiou et al., 2019), the mean BOLD fMRI signal within larger regions of interest was initially used to obtain dCVR curves using standard (not spherical) Laguerre expansions (Marmarelis 1993; Marmarelis 2004), and these curves were subsequently used to define the extended gamma set that was used in their analyses. In turn, this suggests that such an initialization step should be repeated for different data sets. Indeed, as the BEG method does not use adaptive basis sets, its basis set should ideally be tuned for each type of data and/or application.

The CCA method also uses a Gamma basis, but instead of simplifying the basis set through variance and PCA (BEG method), it does so in the temporal dimension by including derivative terms (Friston et al., 1998; Henson et al., 2002; Hossein-Zadeh et al., 2003a). These derivatives allow for estimated *h*(*t*) to be shifted in either direction in time, thus allowing for temporal features of dCVR to be prioritized. Nonetheless, though applied with some success in estimating the neuronal hemodynamic response (Hossein-Zadeh et al., 2003a), it performed the worst for dCVR estimation, being the most variable and least accurate. In this regard, it also displayed minimal SNR dependence. One reason for this is that the conventional CCA method adopts a fixed set of model parameters instead of being adaptive, and is incapable of representing *h*(*t*) in high or low-SNR conditions despite its use of temporal derivatives. Thus, the conventional CCA method is included as an example of simplicity and inflexibility for the application at hand. In contrast, the CCA_opt_ method leverages the simplicity of the conventional CCA method, but with the proposed optimization that allows the parameters of the model basis to vary.

### Ground-truth dependence of modeling methods

In practice, it is impossible to know the underlying shape of *h*(*t*). Although the shape of *h*(*t*) in the healthy brain has been informed by the many works demonstrating the hemodynamic response function (Aguirre et al. 1998; Shan et al. 2014), different disease conditions have been known to alter the dCVR shape. For instance, aging (West et al., 2019), autism spectrum disorder (Yan et al., 2018) and Alzheimer’s disease (Morsheddost et al., 2014) have all been known to alter the response shape. Thus, by evaluating all methods on a wide range of HRF timings and shapes, we can determine the suitability of these model-based methods for estimating an unknown *h*(*t*), one that may not conform to any of the four models.

The performances of these model-based methods were ground-truth dependent, but to our surprise, the performance of the methods was not dictated by their underlying model assumptions. The poor overall performance of the CCA method, for one, may have been the result of the same limitations discussed in the previous section.

### Computational time

The timing information provides an additional dimension of data for choosing methods. The IL and BEL methods are associated with the most accurate and robust performance, but are also correspondingly computational intensive. The CCA_opt_ method, while ranking just below these two methods in performance, is substantially faster. Specifically, for the IL method, optimal performance required parameter search by the simulated annealing option, thus making it 100 times slower than the optimized CCA method.

### Limitations

In this comparison, we did not include all possible modeling methods, such as the cosine-basis approach (Zarahn 2002), the radial-basis approach (Riera et al., 2004), and the basis-function optimization strategy (Riera et al., 2004; Woolrich et al., 2004). Our rationale is to target methods that are more recent (IL) and also more relevant to dCVR mapping (MAP, BEG, BEL). However, our findings of model-independence can generalize to other methods. Moreover, we provide clear evidence that model-based methods can retrieve the dCVR amidst high noise. We also provide, amongst the methods evaluated, a quantitative basis for choosing a method, based on the desired dCVR parameters, the accuracy and computation time. These findings lay the foundation for wider adoption of dCVR estimation in CVR mapping.

## Acknowledgements

This work has been supported by the Canadian Institutes of Health Research and the Sandra Rotman Foundation.

## References

Aguirre, G.K., Zarahn, E., D’Esposito, M., 1998. The variability of human BOLD hemodynamic responses. NeuroImage, https://doi.org/10.1016/s1053-8119(18)31407-1

Atwi, S., Shao, H., Crane, D.E., da Costa, L., Aviv, R.I., Mikulis, D.J., Black, S.E., MacIntosh, B.J., 2019. BOLD-based cerebrovascular reactivity vascular transfer function isolates amplitude and timing responses to better characterize cerebral small vessel disease. NMR Biomed. e4064.

Battisti-Charbonney, A., Fisher, J., Duffin, J., 2011. The cerebrovascular response to carbon dioxide in humans. J physiology 589, 3039–3048.

Blockley, N.P., Harkin, J.W., Bulte, D.P., 2017. Rapid cerebrovascular reactivity mapping: Enabling vascular reactivity information to be routinely acquired. Neuroimage. https://doi.org/10.1016.j.neuroimage.2017.07.048

Chang, C., Cunningham, J.P., Glover, G.H., 2009. Influence of heart rate on the BOLD signal: the cardiac response function. Neuroimage 44, 857–869.

Chang, C., Glover, G.H., 2009. Relationship between respiration, end-tidal CO2, and BOLD signals in resting-state fMRI. Neuroimage 47, 1381–1393.

Chen, J.J., 2018. Cerebrovascular-Reactivity Mapping Using MRI: Considerations for Alzheimer’s Disease. Front. Aging Neurosci.

Chen, J.J., Golestani, A.M., Wei, L., 2021. Quantitative mapping of cerebrovascular reactivity using resting-state functional magnetic resonance imaging. US Patent 10,898,143.

Chen, J.J., Smith, M.R., Frayne, R., 2005. Advantages of frequency-domain modeling in dynamic-susceptibility contrast magnetic resonance cerebral blood flow quantification. Magn. Reson. Med. 53, 700–707.

Chen, K., Yang, H., Zhang, H., Meng, C., Becker, B., Biswal, B., 2021. Altered cerebrovascular reactivity due to respiratory rate and breath holding: a BOLD-fMRI study on healthy adults. Brain Struct. Funct. https://doi.org/10.1007/s00429-021-02236-5

Dabir, A.S., Trivedi, C.A., Ryu, Y., Pande, P., Jo, J.A., 2009. Fully automated deconvolution method for online analysis of time-resolved fluorescence spectroscopy data based on an iterative Laguerre expansion technique. Journal of Biomedical Optics. https://doi.org/10.1117/1.3103342

Davis, T.L., Kwong, K.K., Weisskoff, R.M., Rosen, B.R., 1998. Calibrated functional MRI: mapping the dynamics of oxidative metabolism. Proc. Natl. Acad. Sci. U. S. A. 95, 1834–1839.

Duffin, J., Sobczyk, O., Crawley, A.P., Poublanc, J., Mikulis, D.J., Fisher, J.A., 2015. The dynamics of cerebrovascular reactivity shown with transfer function analysis. Neuroimage 114, 207–216.

Fierstra, J., Sobczyk, O., Battisti-Charbonney, A., Mandell, D.M., Poublanc, J., Crawley, A.P., Mikulis, D.J., Duffin, J., Fisher, J.A., 2013. Measuring cerebrovascular reactivity: what stimulus to use? J. Physiol. 591, 5809–5821.

Francis, D.P., Seydnejad, S.R., Kitney, R.I., Coats, A.J.S., n.d. Dynamic chemoreceptor response using Laguerre expansion technique. Proceedings of the First Joint BMES/EMBS Conference. 1999 IEEE Engineering in Medicine and Biology 21st Annual Conference and the 1999 Annual Fall Meeting of the Biomedical Engineering Society (Cat. No.99CH37015). https://doi.org/10.1109/iembs.1999.804171

Friston, K.J., Josephs, O., Rees, G., Turner, R., 1998. Nonlinear Event-Related Responses in fMRI. Magn. Reson. Med. 39, 41–52.

Glodzik, L., Randall, C., Rusinek, H., de Leon, M.J., 2013. Cerebrovascular reactivity to carbon dioxide in Alzheimer’s disease. J. Alzheimers. Dis. 35, 427–440.

Glover, G.H., 1999. Deconvolution of Impulse Response in Event-Related BOLD fMRI1. Neuroimage 9, 416–429.

Gokcal, E., Horn, M.J., Becker, J.A., Das, A.S., Schwab, K., Biffi, A., Rost, N., Rosand, J., Viswanathan, A., Polimeni, J.R., Johnson, K.A., Greenberg, S.M., Gurol, M.E., 2022. Effect of vascular amyloid on white matter disease is mediated by vascular dysfunction in cerebral amyloid angiopathy. J. Cereb. Blood Flow Metab. 271678X221076571.

Golestani, A.M., Chang, C., Kwinta, J.B., Khatamian, Y.B., Chen, J.J., 2015. Mapping the end-tidal CO2 response function in the resting-state BOLD fMRI signal: Spatial specificity, test–retest reliability and effect of fMRI sampling rate. Neuroimage 104, 266–277.

Golestani, A.M., Wei, L.L., Chen, J.J., 2016. Quantitative mapping of cerebrovascular reactivity using resting-state BOLD fMRI: Validation in healthy adults. Neuroimage 138, 147–163.

Goutte, C., Nielsen, F.A., Hansen, L.K., 2000. Modeling the haemodynamic response in fMRI using smooth FIR filters. IEEE Trans. Med. Imaging 19, 1188–1201.

Halani, S., Kwinta, J.B., Golestani, A.M., Khatamian, Y.B., Chen, J.J., 2015. Comparing cerebrovascular reactivity measured using BOLD and cerebral blood flow MRI: The effect of basal vascular tension on vasodilatory and vasoconstrictive reactivity. Neuroimage 110, 110–123.

Henson, R.N.A., Price, C.J., Rugg, M.D., Turner, R., Friston, K.J., 2002. Detecting latency differences in event-related BOLD responses: application to words versus nonwords and initial versus repeated face presentations. Neuroimage 15, 83–97.

Hoge, R.D., Atkinson, J., Gill, B., Crelier, G.R., Marrett, S., Pike, G.B., 1999. Investigation of BOLD signal dependence on cerebral blood flow and oxygen consumption: the deoxyhemoglobin dilution model. Magn. Reson. Med. 42, 849–863.

Holmes, K.R., Tang-Wai, D., Sam, K., McKetton, L., Poublanc, J., Crawley, A.P., Sobczyk, O., Cohn, M., Duffin, J., Tartaglia, M.C., Black, S.E., Fisher, J.A., Wasserman, B., Mikulis, D.J., 2020. Slowed Temporal and Parietal Cerebrovascular Response in Patients with Alzheimer’s Disease. Can. J. Neurol. Sci. 47, 366–373.

Hossein-Zadeh, G.-A., Ardekani, B.A., Soltanian-Zadeh, H., 2003a. Activation detection in fMRI using a maximum energy ratio statistic obtained by adaptive spatial filtering. IEEE Trans. Med. Imaging 22, 795–805.

Hossein-Zadeh, G.-A., Ardekani, B.A., Soltanian-Zadeh, H., 2003b. A signal subspace approach for modeling the hemodynamic response function in fMRI. Magnetic Resonance Imaging. https://doi.org/10.1016/s0730-725x(03)00180-2

Ishii, Y., Thamm, T., Guo, J., Khalighi, M.M., Wardak, M., Holley, D., Gandhi, H., Park, J.H., Shen, B., Steinberg, G.K., Chin, F.T., Zaharchuk, G., Fan, A.P., 2020. Simultaneous phase-contrast MRI and PET for noninvasive quantification of cerebral blood flow and reactivity in healthy subjects and patients with cerebrovascular disease. J. Magn. Reson. Imaging 51, 183–194.

Jahanian, H., Christen, T., Moseley, M.E., Pajewski, N.M., Wright, C.B., Tamura, M.K., Zaharchuk, G., SPRINT Study Research Group, 2017. Measuring vascular reactivity with resting-state blood oxygenation level-dependent (BOLD) signal fluctuations: A potential alternative to the breath-holding challenge? J. Cereb. Blood Flow Metab. 37, 2526–2538.

Leistedt, B., McEwen, J.D., 2012. Exact Wavelets on the Ball. IEEE Transactions on Signal Processing. https://doi.org/10.1109/tsp.2012.2215030

Leoni, R.F., Mazzeto-Betti, K.C., Andrade, K.C., de Araujo, D.B., 2008. Quantitative evaluation of hemodynamic response after hypercapnia among different brain territories by fMRI. Neuroimage 41, 1192–1198.

Li, M., Newton, A.T., Anderson, A.W., Ding, Z., Gore, J.C., 2019. Characterization of the hemodynamic response function in white matter tracts for event-related fMRI. Nat. Commun. 10, 1–11.

Lindquist, M.A., Loh, J.M., Lauren Y. Atlas, Wager, T.D., 2009. Modeling the hemodynamic response function in fMRI: Efficiency, bias and mis-modeling. NeuroImage. https://doi.org/10.1016/j.neuroimage.2008.10.065

Lindquist, M.A., Wager, T.D., 2007. Validity and power in hemodynamic response modeling: a comparison study and a new approach. Hum. Brain Mapp. 28, 764–784.

Lindquist, M., Wager, T., 2005. Modeling the hemodynamic response function using inverse logit functions, in: Proceedings of the Human Brain Mapping Annual Meeting.

Liu, P., Li, Y., Pinho, M., Park, D.C., Welch, B.G., Lu, H., 2017. Cerebrovascular reactivity mapping without gas challenges. Neuroimage 146, 320–326.

Lu, Y., Grova, C., Kobayashi, E., Dubeau, F., Gotman, J., 2007. Using voxel-specific hemodynamic response function in EEG-fMRI data analysis: An estimation and detection model. Neuroimage 34, 195–203.

Mandell, D.M., Han, J.S., Poublanc, J., Crawley, A.P., Kassner, A., Fisher, J.A., Mikulis, D.J., 2008. Selective reduction of blood flow to white matter during hypercapnia corresponds with leukoaraiosis. Stroke 39, 1993–1998.

Marmarelis, V.Z., 2004. Nonlinear Dynamic Modeling of Physiological Systems. https://doi.org/10.1002/9780471679370

Marmarelis, V.Z., 1993. Identification of nonlinear biological systems using Laguerre expansions of kernels. Ann. Biomed. Eng. 21, 573–589.

Mayer, A.R., Toulouse, T., Klimaj, S., Ling, J.M., Pena, A., Bellgowan, P.S.F., 2014. Investigating the properties of the hemodynamic response function after mild traumatic brain injury. J. Neurotrauma 31, 189–197.

Morsheddost, H., Asemani, D., Shalchy, M.A., 2014. Effects of aging on BOLD hemodynamic response: Healthy aging versus Alzheimer disease. 2014 22nd Iranian Conference on Electrical Engineering (ICEE). https://doi.org/10.1109/iraniancee.2014.6999852

Nowak-Flück, D., Ainslie, P.N., Bain, A.R., Ahmed, A., Wildfong, K.W., Morris, L.E., Phillips, A.A., Fisher, J.P., 2018. Effect of healthy aging on cerebral blood flow, CO2 reactivity, and neurovascular coupling during exercise. J. Appl. Physiol. 125, 1917–1930.

Ostergaard, L., Weisskoff, R.M., Chesler, D.A., Gyldensted, C., Rosen, B.R., 1996. High resolution measurement of cerebral blood flow using intravascular tracer bolus passages. Part I: Mathematical approach and statistical analysis. Magn. Reson. Med. 36, 715–725.

Peng, S.-L., Chen, X., Li, Y., Rodrigue, K.M., Park, D.C., Lu, H., 2018. Age-related changes in cerebrovascular reactivity and their relationship to cognition: A four-year longitudinal study. Neuroimage 174, 257–262.

Pillai, J.J., Mikulis, D.J., 2015. Cerebrovascular Reactivity Mapping: An Evolving Standard for Clinical Functional Imaging. AJNR Am. J. Neuroradiol. 36, 7–13.

Pinto, J., Bright, M.G., Bulte, D.P., Figueiredo, P., 2020. Cerebrovascular Reactivity Mapping Without Gas Challenges: A Methodological Guide. Front. Physiol. 11, 608475.

Poublanc, J., Crawley, A.P., Sobczyk, O., Montandon, G., Sam, K., Mandell, D.M., Dufort, P., Venkatraghavan, L., Duffin, J., Mikulis, D.J., Fisher, J.A., 2015. Measuring cerebrovascular reactivity: the dynamic response to a step hypercapnic stimulus. J. Cereb. Blood Flow Metab. 35, 1746–1756.

Prokopiou, P.C., Pattinson, K.T.S., Wise, R.G., Mitsis, G.D., 2019. Modeling of dynamic cerebrovascular reactivity to spontaneous and externally induced CO2 fluctuations in the human brain using BOLD-fMRI. Neuroimage 186, 533–548.

Prokopiou, P.C., Xifra-Porxas, A., Kassinopoulos, M., 2020. Modeling the hemodynamic response function using motor task and eyes-open resting-state EEG-fMRI. bioRxiv.

Prokopiou, P.C., Xifra-Porxas, A., Kassinopoulos, M., Boudrias, M.-H., Mitsis, G.D., 2022. Modeling the Hemodynamic Response Function Using EEG-fMRI Data During Eyes-Open Resting-State Conditions and Motor Task Execution. Brain Topogr. https://doi.org/10.1007/s10548-022-00898-w

Rangaprakash, D., Tadayonnejad, R., Deshpande, G., O’Neill, J., Feusner, J.D., 2021. FMRI hemodynamic response function (HRF) as a novel marker of brain function: applications for understanding obsessive-compulsive disorder pathology and treatment response. Brain Imaging Behav. 15, 1622–1640.

Riera, J.J., Watanabe, J., Kazuki, I., Naoki, M., Aubert, E., Ozaki, T., Kawashima, R., 2004. A state-space model of the hemodynamic approach: nonlinear filtering of BOLD signals. NeuroImage. https://doi.org/10.1016/j.neuroimage.2003.09.052

Roustit, M., Blaise, S., Millet, C., Cracowski, J.-L., 2011. Impaired transient vasodilation and increased vasoconstriction to digital local cooling in primary Raynaud’s phenomenon. Am. J. Physiol. Heart Circ. Physiol. 301, H324–30.

Shams, S.-M., Hossein-Zadeh, G.-A., Soltanian-Zadeh, H., 2006. Multisubject activation detection in fMRI by testing correlation of data with a signal subspace. Magn. Reson. Imaging 24, 775–784.

Shan, Z.Y., Wright, M.J., Thompson, P.M., McMahon, K.L., Blokland, G.G.A.M., de Zubicaray, G.I., Martin, N.G., Vinkhuyzen, A.A.E., Reutens, D.C., 2014. Modeling of the hemodynamic responses in block design fMRI studies. J. Cereb. Blood Flow Metab. 34, 316–324.

Sobczyk, O., Battisti-Charbonney, A., Fierstra, J., Mandell, D.M., Poublanc, J., Crawley, A.P., Mikulis, D.J., Duffin, J., Fisher, J.A., 2014. A conceptual model for CO_2_-induced redistribution of cerebral blood flow with experimental confirmation using BOLD MRI. Neuroimage 92, 56–68.

West, K.L., Zuppichini, M.D., Turner, M.P., Sivakolundu, D.K., Zhao, Y., Abdelkarim, D., Spence, J.S., Rypma, B., 2019. BOLD hemodynamic response function changes significantly with healthy aging. Neuroimage 188, 198–207.

Woolrich, M.W., Behrens, T.E.J., Smith, S.M., 2004. Constrained linear basis sets for HRF modelling using Variational Bayes. Neuroimage 21, 1748–1761.

Wu, G.-R., Colenbier, N., Van Den Bossche, S., Clauw, K., Johri, A., Tandon, M., Marinazzo, D., 2021. rsHRF: A toolbox for resting-state HRF estimation and deconvolution. NeuroImage. https://doi.org/10.1016/j.neuroimage.2021.118591

Wu, O., Østergaard, L., Weisskoff, R.M., Benner, T., Rosen, B.R., Sorensen, A.G., 2003. Tracer arrival timing-insensitive technique for estimating flow in MR perfusion-weighted imaging using singular value decomposition with a block-circulant deconvolution matrix. Magn. Reson. Med. 50, 164–174.

Yanai, H., Takane, Y., 1992. Canonical correlation analysis with linear constraints. Linear Algebra and its Applications. https://doi.org/10.1016/0024-3795(92)90211-r

Yan, W., Rangaprakash, D., Deshpande, G., 2018. Estimated hemodynamic response function parameters obtained from resting state BOLD fMRI signals in subjects with autism spectrum disorder and matched healthy subjects. Data in Brief. https://doi.org/10.1016/j.dib.2018.04.126

Zhao, M.Y., Fan, A.P., Chen, D.Y.-T., Sokolska, M.J., Guo, J., Ishii, Y., Shin, D.D., Khalighi, M.M., Holley, D., Halbert, K., Otte, A., Williams, B., Rostami, T., Park, J.-H., Shen, B., Zaharchuk, G., 2021. Cerebrovascular reactivity measurements using simultaneous 150-water PET and ASL MRI: Impacts of arterial transit time, labeling efficiency, and hematocrit. Neuroimage 233, 117955.

