## Supplementary Materials for "Modeling the carbon-dioxide response function in fMRI under task and resting-state conditions"

#### Canonical-correlation analysis (CCA) method

Given two column vectors $Y=\left[ Y^{1} Y^{2} \cdot\cdot\cdot Y^{n} \right]$ and $Ŷ=\left[ Ŷ^{1} Ŷ^{2} \cdot\cdot\cdotŶ^{m} \right]$ of random variables, CCA finds optimal weighting vectors *W_Y_* and $W_{Ŷ}$ such that the correlation between *W_Y_Y* and $W_{Ŷ}Ŷ$*W* is maximized, subject to linear constraints (Yanai and Takane, 1992). It has been shown that the canonical vectors, *W_Y_* and *W_Ŷ_* are the eigenvectors corresponding to the largest eigenvalues of the matrices $S_{Ŷ}^{-1}S_{ŶY}S_{Y}^{-1}S_{YŶ}$ and $S_{Ŷ}^{-1}S_{ŶY}S_{Y}^{-1}S_{YŶ}$, respectively, where $S_{Y}=Y^{T}Y$ and $S_{Ŷ}=Ŷ^{T}Ŷ$ are the within set covariance matrices and $S_{YŶ}=Y^{T}Ŷ$ and $S_{ŶY}=Ŷ^{T}Y=S_{YŶ}^{T}$ are the cross-covariance matrices of the vectors in the Y and $Ŷ$ column vectors. Here we define *Y* as the vector of BOLD signal measurements (BOLD(*t*)), and $Ŷ$ is the column vector of the convolutions between the assumed $h(t)$. $h(t)$ here is assumed to be a single Gamma function

$h\left( t;A,\tau,\sigma\right)=A\cdot\{exp\left( -t/\sqrt{\sigma t} \right)\left( \frac{e.t}{\tau} \right)^{\sqrt{\frac{\tau}{\sigma}}}, t>0$	(A1)

$0, t<0$

where σ represents the width of the peak (dispersion) and τ its location. The temporal derivatives of $h(t)$ and the stimulus pattern (e.g., PETCO_2_(t)), and $\hat{h}(t)$ is the initialization for $h(t)$:

$Ŷ=\left[ ĥ\left( t; \tau, \sigma\right)*X\left( t \right), \frac{\partial}{\partial\tau}ĥ\left( t; \tau, \sigma\right)*X\left( t \right), \frac{\partial}{\partial\sigma}ĥ\left( t; \tau, \sigma\right)*X\left( t \right) \right]$	(A2)

As mentioned, CCA finds the optimal weighting vectors ($W_{Y}$ and $W_{Ŷ}$) that maximize the correlation between column vectors Y and $Ŷ$,

$U=w_{Ŷ}^{1}\cdotĥ\left( t; \tau, \sigma\right)*X\left( t \right)+w_{Ŷ}^{2}\cdot\frac{\partial}{\partial\tau}ĥ\left( t; \tau, \sigma\right)*X\left( t \right)+w_{Ŷ}^{3}\cdot\frac{\partial}{\partial\sigma}ĥ\left( t; \tau, \sigma\right)*X\left( t \right)$	(A3)

$V=w_{Y}\cdot Y(t)$	(A4)
